# The cerebellar clock: predicting and timing somatosensory touch

**DOI:** 10.1101/2020.10.01.321455

**Authors:** Lau M. Andersen, Sarang S. Dalal

**Affiliations:** Center of Functionally Integrative Neuroscience (CFIN), Aarhus University, Nørrebrogade 44, Building 1A, 8000 Aarhus C, Denmark; Aarhus Institute of Advanced Studies (AIAS), Aarhus University, Høegh-Guldbergs Gade 6B, 8000 Aarhus C, Denmark; National Facility for Magnetoencephalography (NatMEG), Karolinska Institutet, Nobels väg 9, 171 77 Stockholm, Sweden

## Abstract

The prediction of sensory feedback is known to involve the cerebellum, but its precise nature and timing have remained unclear due to the scarcity of time-sensitive cerebellar neuroimaging studies. We here, using magnetoencephalography in human participants, investigated the working hypothesis that one function of the cerebellum is to predict exactly when rhythmic stimuli are expected to impinge on sensory receptors.

We compared the cerebellum’s response between somatosensory omissions embedded in perfectly rhythmic versus jittered trains of stimulation. At the precise moment that an omitted stimulus would have appeared, the cerebellum exhibited a beta band (14-30 Hz) response that was stronger when preceded by a perfectly rhythmic sequence. Meanwhile, the rhythm of new stimulation sequences induced theta band (4-7 Hz) activity in the cerebellum.

Our results provide evidence that the cerebellum acts as a clock that entrains to rhythmic stimuli, likely for the purpose of detecting any deviations from that rhythm.

## 1 Introduction

It has become clear that the cerebellum actively predicts what feedback it should receive from the external world (Hull, 2020). In this study, we investigate the working hypothesis that the cerebellum functions as a clock, tracking exactly when somatosensory stimuli are expected to impinge on sensory receptors. Direct evidence of sub-second sensory prediction has been found in the monkey (Ohmae et al., 2013), but neural evidence from humans is scarce. That the cerebellum functions as a clock is in line with older proposals suggesting, for example, that the fundamental purpose of the cerebellum is to predict and set conditions necessary for motor and mental operations (Courchesne and Allen, 1997), or that the cerebellum builds a forward model, which sensory input can be checked against, (Ramnani, 2006). Furthermore, patients with cerebellar lesions reveal difficulties in tasks that require processing of temporal information (Schwartze et al., 2016). The objective of the current study was to test the timing and precision of cerebellar responses in the millisecond range. We measured the cerebellum’s response to omitted somatosensory stimuli in stimulation trains of varying regularity, using the non-invasive, time-sensitive method of magnetoencephalography (MEG). Tesche and Karhu (2000) provided early MEG evidence suggesting that the cerebellum updates expectations. Estimating time courses for dipolar sources in the cerebellar vermis, they found oscillatory responses to omitted somatosensory stimuli in the ranges 6-12 Hz and 25-35 Hz. These were maximal around the time the stimulus should have happened, indicating a relationship to expectational processes. We, (Andersen and Lundqvist, 2019), found direct evidence that the cerebellum was more strongly activated in the first repetition of a stimulus compared to the first stimulation for the ranges 4-7 Hz and 10-30 Hz. Furthermore, we found a tonic difference between omitted stimulations, which are unexpected absences of stimulation, and non-stimulations, which are expected absences of stimulation, in the range of 3-15 Hz. This may be an effect of more power in the mu-rhythm when the body is at rest (Kuhlman, 1978), as is the case for non-stimulations. Thus, to find a time-dependent cerebellar response to omissions, we reasoned that omissions of different types should be contrasted against one another instead of against non-stimulations.

The main hypothesis of the current study is that the cerebellum will be more strongly activated to omissions of otherwise expected stimulation during regular than during irregular stimulation trains. We expect this difference in cerebellar activation to occur around the expected timing of the omitted stimulus. Based on the findings in Andersen and Lundqvist (2019) and Tesche and Karhu (2000) we expect these differences to occur in the theta (4-7 Hz) and beta (14-30 Hz) bands. Besides the cerebellum, we investigated other areas implicated in somatosensation and in updating sensory expectations, specifically the primary and secondary somatosensory cortices (SI and SII), and the inferior parietal cortex (Andersen and Lundqvist, 2019; Tesche and Karhu, 2000).

The cerebellum has had a reputation of being impossible to study with non-invasive electrophysiological methods due to it being finely folded, which should result in signal cancellation. However, we, (Andersen et al., 2020), recently reviewed the magneto-and electroencephalographic literature on cerebellar findings and found at least thirty studies reporting cerebellar activity. Supporting the validity of these findings, Samuelsson et al. (2020) recently showed, using a combination of MEG signal simulation and high-resolution magnetic resonance imaging (MRI), capturing the fine structure of the cerebellum, that cerebellar signals are strong enough to be detectable by state-of-the-art MEG. Studies on the turtle and pig brains have also shown that the cerebellum produces magnetic fields measurable at a distance (Okada et al., 1997, 1987).

It has been proposed that the basal ganglia and the cerebellum together with the thalamus are nodes of an integrated network responsible for cognitive timing (Bostan and Strick, 2018; Gibbon et al., 1997). Due to both their depth and the possibility that their neurons may form a closed field arrangement (Lorente de Nó, 1947), MEG has traditionally not been considered sensitive to these structures. However, Attal et al. (2012) argue that MEG should be sensitive to the anterior parts of the putamen, showing that it should contribute to ∼10% of the total MEG signals. Attal and Schwartz (2013) also showed experimental evidence that an increase in thalamic alpha band power when subjects had their eyes closed could be measured with MEG. Pizzo et al. (2019) recently showed that time courses of thalamic activation measured by stereotactic electroencephalography correlated with MEG. Therefore, we also included the putamen and the thalamus in our MEG source analyses in order to examine their potential role in a cognitive timing network.

## 2 Materials and Methods

### 2.1 Participants

Thirty right-handed, healthy participants volunteered to take part in the experiment (seventeen males and thirteen females, Mean Age: 26.7 y; Standard Deviation: 6.3 y; Range: 18-41 y). The experiment was advertised in a local database that participants had signed up to. The experiment was approved by the local institutional review board in accordance with the Declaration of Helsinki. The participants provided informed consent for participation and were compensated by 100 DKK per hour (on average 3 hours of experimentation over two sessions).

### 2.2 Stimuli and procedure

Tactile stimulation was generated by two ring electrodes driven by an electric current generator (Stimulus Current Generator, DeMeTec GmbH, Germany). The ring electrodes were fastened to the tip of the right index finger. One was placed 10 mm below the bottom of the nail and the other 10 mm below that. Stimuli were applied in sequences of six stimuli, with an inter-stimulus interval of 1,497 ms, such that the stimulation would not lock to the 50 Hz power from electrics. All pulses had a duration of 100 µs, and for each participant an individualised level of current was found as described below. The sixth stimulus was followed by an omission, and a new train of stimulation began. Thus, between the last stimulus of a train and the following train 2,994 ms elapsed. Three types of stimulation sequences and one type of non-stimulation sequence were administered (Fig. 1). 1) a no-jitter sequence, where all stimuli happened exactly on time (0 % jitter); 2) a jittered sequence, where stimuli 4-6 were 5 % jittered, i.e. they happened from ∼-75 ms to ∼75 ms (in integer steps of 1 ms) relative to where the stimulus would have happened, had the sequence been steady; 3) a heavily jittered sequence, where stimuli 4-6 were 15 % jittered, i.e. they happened from ∼-225 ms to ∼225 ms (in integer steps of 1 ms) relative to where the stimulus would have happened, had the sequence been steady. For each jittered stimulation, the jitter, as an integer number of milliseconds, was chosen randomly from a uniform distribution, with the qualification that the stimulation had to be a minimum of 13 ms away from the time where the steady stimulus would have happened. After every fifteen sequences, 4) a train of five non-stimulations would occur, i.e. 5 x 1,497 ms of no stimulation. Thus, omissions and non-stimulations differed by whether or not an expectation was present in the time leading up to it.

**Fig 1.**
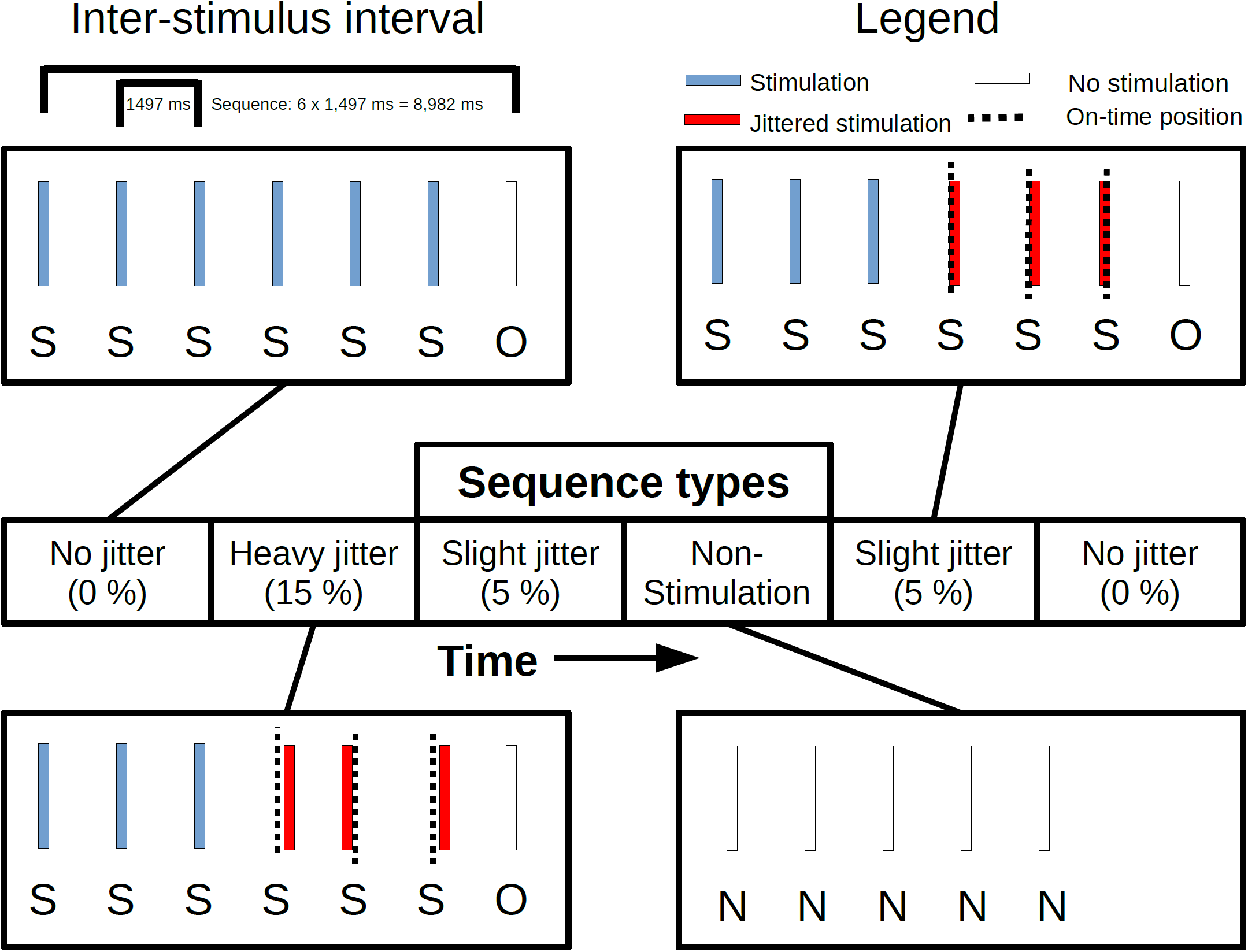
Six instances of four sequence types following one another. The stimuli in the sequences were presented with an inter-stimulus interval of 1,497 ms: The four sequences were: 1) A 0%-jitter type where all stimulations were on-time. 2) A 5%-jitter type where the first three stimuli (blue bars) were on time while the last three stimuli (red bars) were up to 5% off-time, 3) A 15%-jitter type, following the same structure, where the jittered stimuli were up to 15 % off-time. 4) A no-stimulation type, where nothing happened for five times the inter-stimulus interval. The omissions and non-stimulation (white bars) were timed to where the stimulus would have occurred, had it happened and had it been on-time. Dashed vertical lines indicate where the jittered stimulus should have occurred, had it been on time. Abbreviations: S = Stimulation O = Omission, N = Non-Stimulation.

150 stimulation sequences of each type and 30 non-stimulation sequences were administered. The sequences were pseudorandomly interleaved in a counterbalanced manner. This structure resulted in 450 *First Stimulations*, 450 *Second Stimulations* and 450 *Third Stimulations*. Note that the first three stimulations are identical in all three stimulation sequence types and are thus collapsed across the three types. Furthermore, this resulted in 3 x 150, one for each of the three stimulation sequence types, *Fourth Stimulations, Fifth Stimulations* and *Sixth Stimulations*. These last three conditions were not subsequently tested. Finally, it resulted in 3 x 150 *Omitted Stimulations* (*Omission 0, Omission 5* and *Omission 15*) and a total of 150 *Non-Stimulations*. Six breaks, with equally long sessions between them were administered throughout, where verbal contact was initiated between the experimenter and the participant to check whether everything was alright. PsychoPy (version 3.2.4) (Peirce, 2009) was used to deliver the stimuli from a Linux operating system running Ubuntu Mate (kernel version 4.15.0-126).

During the stimulation procedure, participants watched a nature programme with sound from panel speakers. Participants were instructed to pay full attention to the movie, focus on the centre of the screen, stay motionless and pay no attention to the stimulation of the finger.

Electro-oculography, -cardiography and myography (EOG, ECG and EMG) were recorded. For EOG, eye movements and eye blinks were recorded to monitor that participants did not blink excessively or moved their eyes away from the centre of the screen. EMG was recorded over the splenius muscles. These were used to monitor that participants would not build up tension in the neck muscles. Lastly, for explorative purposes, respiration was measured using a respiratory belt but is not analysed here.

### 2.3 Preparation of participants

In preparation for the measurement, each participant had two pairs of EOG electrodes placed horizontally and vertically, respectively, around the eyes. ECG was measured by having a pair of electrodes on each collarbone. Finally, two pairs of EMG electrodes were placed on either side of the splenius muscles. Four head-position indicator (HPI) coils, two behind the ears and two on the forehead, were placed on the participants. The ground electrode was placed on the right arm above the ring electrode. Subsequently, each participant had their head shape digitized using a Polhemus FASTRAK. Three fiducial points, the nasion and the left and right pre-auricular points, were digitized along with the positions of the HPI-coils. Furthermore, around 200 extra points digitizing the head shape of each participant were acquired. Participants were subsequently placed in the supine position of the MEG system and great care was taken, such that they would lie comfortably in the scanner, thus preventing neck tension. The ring electrodes were then fastened to the tip of the right index finger. One was placed 10 mm below the bottom of the nail and the other 10 mm below that. Before beginning the actual experiment, a participant specific stimulation current was found. The initial current was 1.0 mA. The current was increased in steps of 0.5 mA until the participant reported feeling the current. After that, current was increased in steps of 0.2 mA, until the participant reported it to be irritating. Then, in steps of 0.2 mA, the current was decreased, until it was no longer irritating. It was emphasized to participants that the stimulation should be clear, but neither irritating nor painful. The individual current reached was then used throughout the experiment.

### 2.4 Acquisition of data

Data was recorded on an Elekta Neuromag TRIUX system inside a magnetically shielded room (Vacuumschmelze Ak3b) at a sampling frequency of 1,000 Hz. As data was acquired online, low-pass and high-pass filters were applied, at 330 Hz and 0.1 Hz respectively.

### 2.5 Processing of MEG data

We analysed evoked responses and oscillatory responses. MaxFilter was not applied to preserve the rank of the data. The four first signal space projections (SSPs) were projected out for the sensor space analyses due to the strong influence of gradients on the magnetometers. Sensors were inspected by investigating the power spectral densities of the raw data, but no sensors were removed as none appeared bad, i.e. having power much higher or lower than the average across all frequencies.

All MEG analyses were carried out in MNE-Python (Gramfort et al., 2013b). For the evoked responses, we low-pass filtered the data at 40 Hz (one-pass, non-causal; finite impulse response; zero-phase; upper transition bandwidth: 10.00 (−6 dB cutoff frequency: 45 Hz; filter length: 331 samples; a 0.0194 passband ripple; and 53 dB stopband attenuation) and then cut the raw data into epochs of 800 ms, 200 ms pre-stimulus and 600 ms post-stimulus. The epochs were demeaned using the mean value of the pre-stimulus period. Segments of data including magnetometer responses greater than 4 pT were rejected. Subsequently, epochs were averaged to create evoked responses for each of the trial types. For each of the stimulation conditions, this resulted in a mean of 442 segments per subject across condition. For each of the omission conditions and the non-stimulation condition, this resulted in a mean of 147 segments per subject across condition. Evoked responses for individual participants were inspected and no obvious bad channels were detected before a grand average over all participants was calculated.

For the oscillatory responses, the data were band-pass filtered into the theta and beta bands, both one-pass, non-causal; finite impulse response; zero-phase with a -6 dB cutoff frequency with a 0.0194 passband ripple; and 53 dB stopband attenuation. The bands were defined as from 4-7 Hz (Theta) and 14-30 Hz (Beta). The lower and upper transition bandwidths for the theta band were 2.00 Hz, resulting in −6 dB cutoff frequencies of 3.00 Hz and 8.00 Hz respectively and a filter length of 1651 samples. The lower and upper transition bandwidths for the beta band were 3.50 Hz and 7.50 Hz respectively, resulting in −6 dB cutoff frequencies of 12.25 Hz and 33.75 Hz respectively and a filter length of 943 samples. A Hilbert transform was applied to both bands and the data were cut into epochs of 1,500 ms, 400 ms pre-stimulus and 1,100 ms post-stimulus for the stimulations and 750 ms pre-stimulus and 750 ms post-stimulus for the omissions. The stimulation epochs were demeaned using the mean value of the pre-stimulus interval up to −50 ms, whereas the omission epochs were demeaned using the whole time-interval. Segments of data including magnetometer responses greater than 4 pT or gradiometer responses greater than 400 pT/m were rejected. For the stimulations in the theta range this resulted in a mean of 440 segments remaining per subject across condition, and in the beta range 450 segments. For the omissions in the theta range this resulted in a mean of 147 segments remaining per subject across condition, and in the beta range 150 segments. Epochs were not rejected based on EOG and EMG due to beamformers (see below) being good at suppressing the artefacts arising from eye blinks and movements. We calculated the envelopes of the Hilbert transformed data. Then, for each contrast of interest, we converted that contrast to a *z*-score, a normal distribution, approximated using a Wilcoxon signed-rank test entering each condition of the contrast into the test. This conversion minimises the effects of outliers since the density of values of the normal distribution asymptotically goes towards zero, the more extreme the value.

### 2.6 Source reconstruction

For both the evoked responses and the oscillatory responses, a linearly constrained minimum variance (LCMV) beamformer (van Veen et al., 1997) was applied. For the former, it was applied to evoked responses and for the latter to the Hilbert transformed epochs.

We acquired sagittal T1 weighted 3D images for each participant using a Siemens Magnetom Prisma 3T MRI. The pulse sequence parameters were: 1 mm isotropic resolution; field of view: 256 mm x 256 mm; 192 slices; slice thickness: 1 mm; bandwidth per pixel: 290 Hz/pixel; flip angle: 9°, inversion time (TI): 1,100 ms; echo time (TE): 2.61 ms; repetition time (TR): 2,300 ms. Based on these images, we did a full segmentation of the head and the brain using FreeSurfer (Dale et al., 1999; Fischl et al., 1999). Subsequently we delineated the head and brain surfaces using the watershed algorithm from MNE-C (Gramfort et al., 2013a). We created a volumetric source space from the brain surface with sources 7.5 mm apart from one another (∼4000 sources). Single compartment boundary element method (BEM) solutions (volume conduction models) were calculated from the brain surfaces. For each participant, the T1 was registered to the participant’s head shape with the fiducials and head shape points acquired with Polhemus FASTRAK. Finally, with the co-registered T1, we created two forward models based on the volume conduction model, the positions of MEG sensors, and the volumetric source space.

For estimating source time courses, the data covariance matrix was estimated based on the post-stimulus period (from 0 ms to 600 ms) for the evoked responses. For the oscillatory responses, the data covariance matrix was estimated based on the post-stimulus time period (from 0 ms to 1,100 ms) for the stimulations, and for the omissions the data covariance matrix was estimated based on the whole time period (−750 ms to 750 ms). A beamformer with the unit-noise-gain constraint was used to compute the spatial filter weights (Sekihara and Nagarajan, 2008). No regularisation was applied to the covariance matrix when estimating the filter weights, since the matrices were well-conditioned. For the filters, we chose the source orientation that would maximise source output. Finally, for the oscillatory responses, we took the absolute value of the complex-valued LCMV-source-time-courses, and averaged over all the epochs to acquire an averaged source time course for both bands (4-7 Hz and 14-30 Hz). For the evoked responses, we also used the absolute value of the estimated source time courses.

For all beamformer computations, only magnetometers were used since these are the sensors most sensitive to deep sources. These source reconstructions were morphed onto *fsaverage*, a common template from FreeSurfer. Note, that signal space projection vectors were not applied on the data to be beamformed since beamformers suppress low-rank external noise well (Sekihara and Nagarajan, 2008).

To investigate contrasts in an unbiased manner, the spatial filter of the LCMV beamformer was estimated based on the covariance of the combined data. Subsequently, this filter was used to estimate the power for each condition in the contrast and finally the contrast for the oscillatory responses was calculated as: (*condition_1 – condition_2*) / (*condition_1 + condition_2*), meaning that the contrast expresses the difference in power as a percentage of total power in the two conditions. For the evoked responses, we looked at the difference (*condition_1 – condition_2*).

### 2.7 Statistical analysis

For the oscillatory responses, we focused on the comparisons between the *Omissions*, with the difference between *Omission 0* and *Omission 15* (No-jitter versus Heavy jitter, Fig. 1) expected to be the strongest. Also the comparison between *First Stimulation* and *Second Stimulation* was interesting due to the difference in expectation and our earlier findings of a difference in the theta band (Andersen and Lundqvist, 2019). We used a cluster permutation test (Maris and Oostenveld, 2007) to address the multiple comparisons problem arising from doing all-sensor and whole-brain analyses. The analyses done in sensor space were done on the whole time range (−400 ms to 1,100 ms for the stimulations and −750 ms to 750 ms for the omission). We used the sensor space analyses to inform the time range of our subsequent tests for the whole-brain analyses, restricting the time range by judging the peak of the response and adding 400 ms in both the negative and positive directions. Whole brain analyses were only run, if the null hypothesis of the corresponding sensor space analysis could be rejected. The null hypothesis of such a test is that data are exchangeable, meaning colloquially speaking that our labelling does not matter. The alternative hypothesis then is that the labelling does matter. The procedure was as follows. First, we conducted a standard *t*-test, calculated using the mean and the standard error across subjects, for each of the time-sensor or time-source combinations (e.g. for the beamformer: 801 time samples and 4,342 sources; > 3 million tests). Then spatio-temporal clusters were formed based on connecting neighbouring sensors/sources and neighbouring time points. Only time-sensor/source combinations that were significant at α = 0.05 were included in clusters. 1024 permutations were then run where the condition labels, e.g. *Omission 0* and *Omission 15*, were shuffled between trials. Using the same procedure as for the correctly labelled trials, spatio-temporal clusters were formed from the significant tests for each of the permutations. A null hypothesis distribution was sampled by adding the value of the largest cluster to the distribution for each permutation. The largest cluster in each permutation was defined as the one that had the largest sum of absolute *t*-values. For each cluster in the correctly labelled trials, the likelihood of that cluster, or one more extreme, was found, its *p*-value, given the sampled null hypothesis distribution. The null hypothesis that the data is exchangeable was then rejected if the largest cluster in the correctly labelled data was associated with a *p*-value that is lower than the alpha value, which we set at 0.05.

If the cluster permutation analyses revealed that the null hypothesis of exchangeability could be rejected, we focused subsequent analyses on the regions discussed in the introduction, i.e. the cerebellum, SI, SII, inferior parietal cortex, the thalamus and the putamen. Other regions were not investigated in more detail, as cluster permutation analyses do not allow a strong inference as to the exact location and timing of the effects (Maris and Oostenveld, 2007). However, the nature of the clusters can still be cast in descriptive terms, and as the areas above have all been implicated in sensory and timing processing, it is of high relevance to describe the timing and activity of source currents in these areas (Sassenhagen and Draschkow, 2019).

For the evoked responses, we did the cluster permutation tests on the source time courses directly, since the SI and SII responses were expected. We focused on the comparisons between *First Stimulation* and *Second Stimulation* and between *Omitted Stimulation* and *Non-Stimulation* following Andersen and Lundqvist (2019). We tested on the full time range (−200 ms to 600 ms).

## 3 Results

### 3.1 Evoked responses

The electric stimulations gave rise to the expected SI and SII activations (Fig. 2)

**Fig 2.**
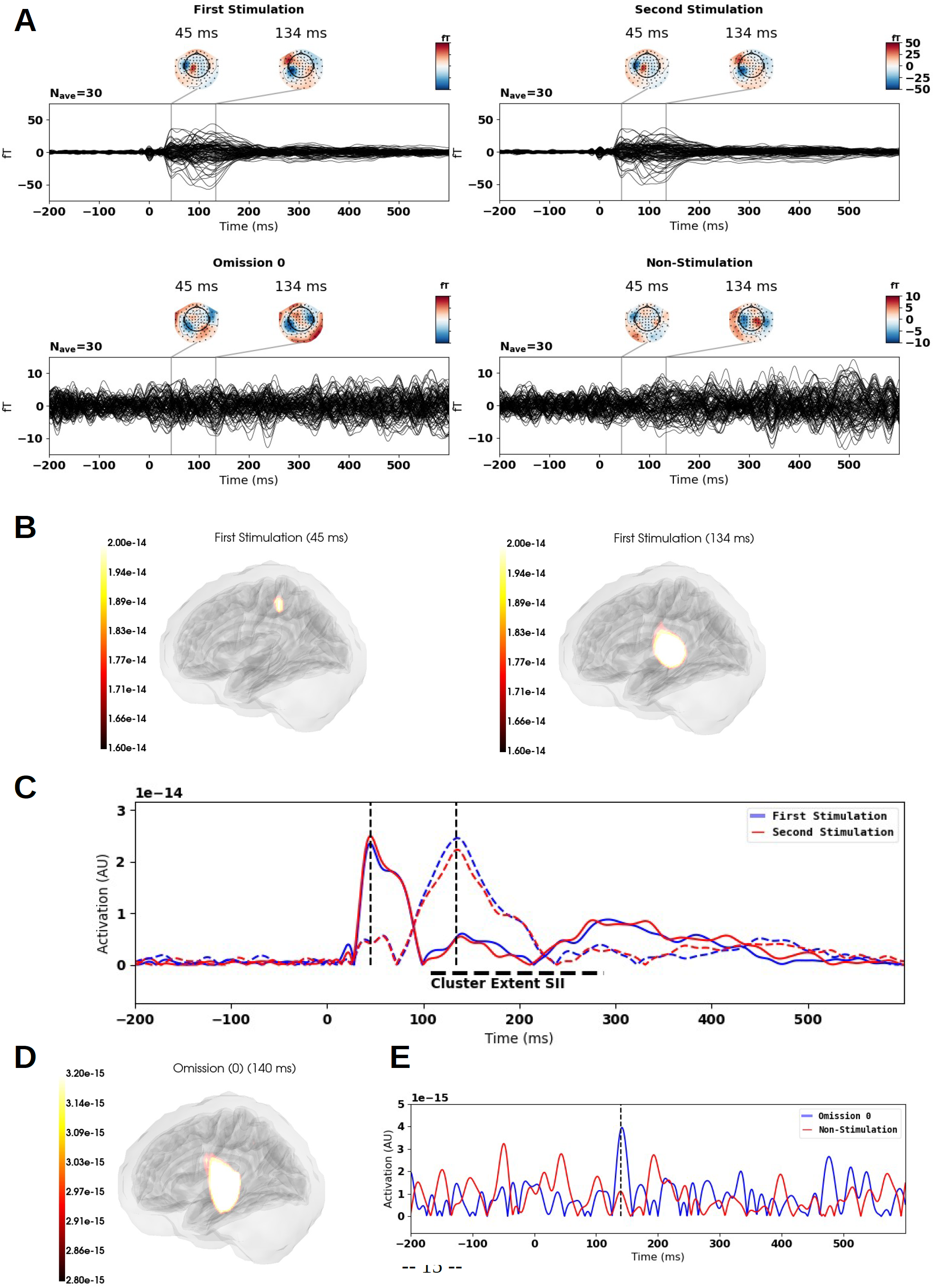
Evoked time courses and source reconstructions. A) Evoked time courses for First Stimulation, Second Stimulation, Omission (0) and Non-Stimulation. The former two show SI and SII time courses, whereas only a weak response is found for the SII response in Omission (0). **B)** Source maps showcasing the early SI activation (left) (45 ms) and joined by the later SII activation (right) (130 ms). **C)** Time courses for the two sources shown in **B**; full lines show SI sources; coloured dashed lines show SII sources; vertical black dashed lines indicate the time points show in **B**. The horizontal black dashed line represents the extremes of the cluster extent for the difference between *First Stimulation* and *Second Stimulation*. **D;** The left SII evoked response for the omission at 140 ms **E;** Evoked time courses for the Omission (0) and Non-Stimulation for the maximally responding source in SII extracted using the Harvard Oxford cortical atlas (Desikan et al., 2006).

An omission response was found (Fig. 2A & 2DE). Differences were found between *First Stimulation* and *Second Stimulation* for the source time courses when using the cluster permutation procedure described above in the time range −200 ms to 600 ms (*p*_BIGGEST_CLUSTER_ = 0.00098). No differences were found between *Omission 0* and *Non-Stimulation* (*p*_BIGGEST_CLUSTER_ = 0.66) using the cluster test. Testing the maximally responding source (Fig. 2E) (140 ms) against 0 does result in a significant test, *t*_29_ = 3.37, *p*_UNCORRECTED_ = 0.002.

### 3.2 Oscillatory responses

For the oscillatory responses, we focused on the theta and beta band responses, following the results of Andersen and Lundqvist (2019) and Tesche and Karhu (2000). For the stimulations, the main contrast of interest was between *First Stimulation* and *Second Stimulation*. In the beta band, the null hypothesis could not be rejected (*p*_BIGGEST_CLUSTER_ = 0.72). In the theta band, we found that 47 magnetometers formed part of two spatiotemporal clusters with *p-*values < 0.05 (*p*_BIGGEST_CLUSTER_ = 0.00098) (Fig. 3A). Given that the responses peaked around 200 ms, we centred the subsequent source cluster analysis on the window of −200 ms to 600 ms. We found one cluster with a *p*-value < 0.05 (*p*_BIGGEST_CLUSTER_ = 0.00098). The maximal value was found for the left inferior parietal cortex at 145 ms (Fig. 3D). A left cerebellar activation was also found after 290 ms (Fig. 3C). Time courses for the contrasts for these two regions were extracted by finding the coordinate that showed the highest contrast (Fig. 3EF). For the comparison between *Second Stimulation* and *Third Stimulation* the null hypothesis could not be rejected (*p*_BIGGEST_CLUSTER_ = 0.12). For descriptive purposes, we reconstructed all the first three stimulations based on a common spatial filter and estimated time courses for the sources shown in Fig. 3EF. These show the cerebellar and inferior parietal cortical activity for each of the conditions in separation. See Supplementary Table 2 for a full list of regions for which more than 50 % of the voxels are part of a cluster.

**Fig 3.**
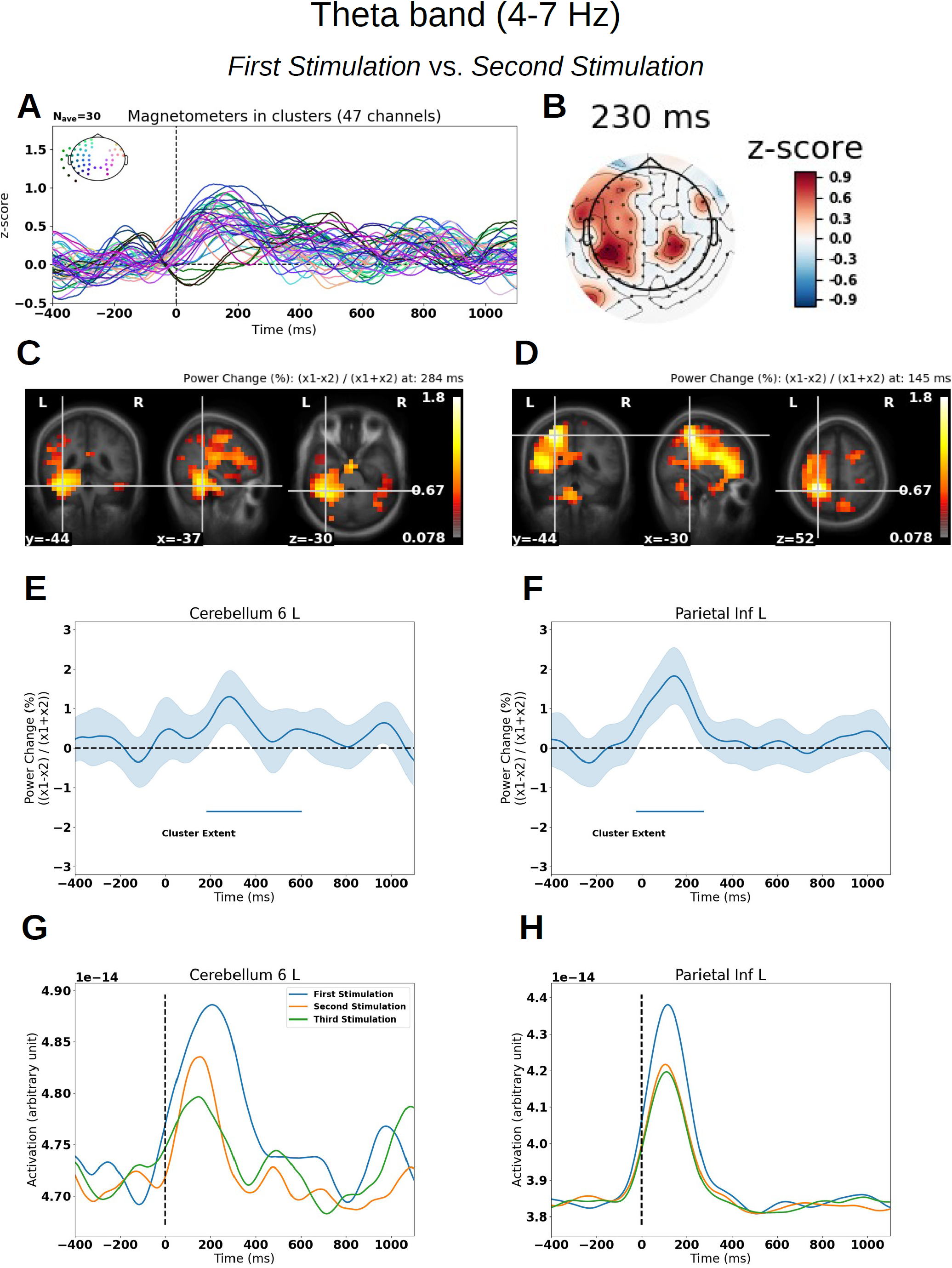
Differences between *First Stimulation* and *Second Stimulation* in the theta band (4-7 Hz) (−200 ms to 600 ms). For all contrast plots, positive values imply greater amplitude in *First Stimulation* than in *Second Stimulation*. **A)** Grand average butterfly plot with colour coded magnetometers according to helmet position. Only magnetometers being part of clusters with a probability less than 0.05 under the sampled null hypothesis distribution are shown **B)** Topographical plot at 230 ms after stimulation, with sensors not forming part of a cluster masked out. **C)** Contrast localized to the left cerebellum at 284 ms: The heat map is thresholded such that values that are not part of a cluster have been set to 0. **D)** The maximum contrast over the whole time course localized to the left inferior parietal cortex at 145 ms: The heat map is thresholded such that values that are not part of a cluster have been set to 0. **E)** and **F)** Time courses for the contrast between *First Stimulation* and *Second Stimulation* with 95% confidence intervals across participants, extracted from the sources in **C** and **D. G)** and **H)** Time courses for *First Stimulation, Second Stimulation* and *Third Stimulation (3)*, extracted from the sources in **C** and **D**. Time courses in **E-H** are extracted from the source showing the maximum activation according to the automated anatomical labeling atlas (AAL) (Tzourio-Mazoyer et al., 2002).

For the omissions, the main contrast of interest was between *Omission 0* and *Omission 15*, as we expected the cerebellar contrast to be the strongest there. In the theta band, the null hypothesis could not be rejected (*p*_BIGGEST_CLUSTER_ = 0.25). In the beta band, we found that 14 magnetometers formed part of three spatiotemporal clusters with *p*-values < 0.05 (*p*_BIGGEST_CLUSTER_ = 0.014) (Fig. 4A). The sensor topography (Fig. 4B) was compatible with an underlying cerebellar source on the right side. For the source reconstruction, we found one cluster with a *p*-value < 0.05 (*p*_BIGGEST_CLUSTER_= 0.0078) testing on the time interval from −400 ms to 400 ms. A right cerebellar source was part of the cluster (Fig. 4C), and the maximally responding source at 0 ms was in the putamen (Fig. 4D).Running similar tests for *Omission 0* vs *Omission 5* and *Omission 5* vs *Omission 15* resulted, however, in not being able to reject the null hypothesis (*p*_BIGGEST_CLUSTER_ = 0.18 and *p*_BIGGEST_CLUSTER_ = 0.11 respectively). Time courses for the contrasts for these two regions were extracted by finding the coordinate that showed the highest contrast (Fig. 4EF). Again, for descriptive purposes, we reconstructed all three types of omissions based on a common spatial filter and estimated time courses for the sources shown in Fig. 4EF. These show that cerebellar activity and putamen activity start increasing around −250 ms of stimulation when the omission is preceded by a regular train of stimulation (*Omission 0*) and starts decreasing again around 0 ms. The opposite pattern emerges for the omissions preceded by irregular trains of stimulation (*Omission 5* and *Omission 15*), which decrease from −250 ms and start increasing from 0 ms. See Supplementary Table 1 for a full list of regions for which more than 50 % of the voxels are part of a cluster.

**Fig 4.**
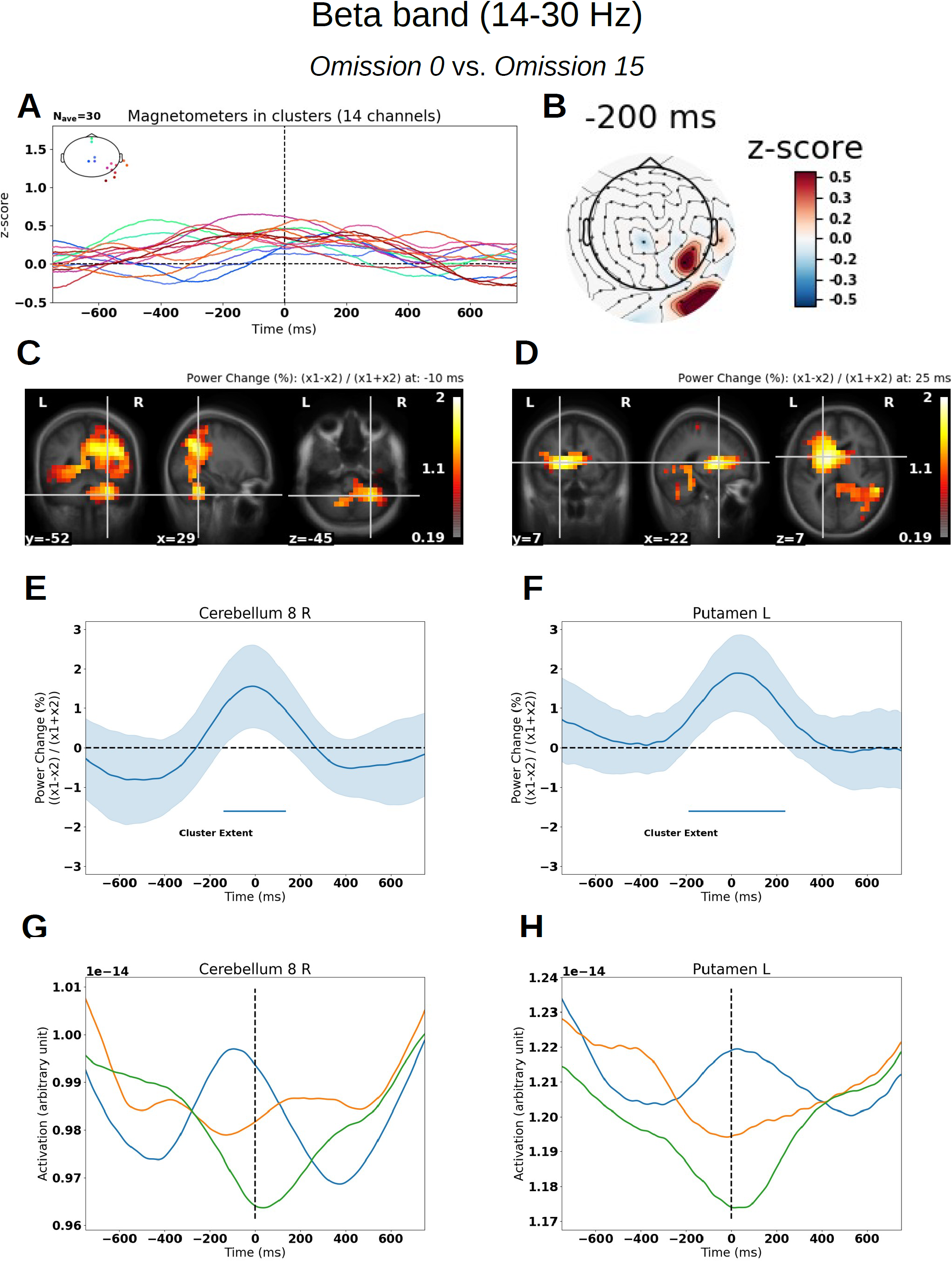
Differences between *Omission 0* and *Omission 15* in the beta band (14-30 Hz) (−400 ms to 400 ms). For all contrast plots, positive values imply greater amplitude in *Omission 0* than in *Omission 15*. **A)** Grand average butterfly plot with colour coded magnetometers according to helmet position. Only magnetometers being part of clusters with a probability less than 0.05 under the sampled null hypothesis distribution are shown. **B)** Topographical plot at −200 ms, with sensors not forming part of a cluster masked out. **C)** Contrast localized to the right cerebellum at −11 ms: The heat map is thresholded such that values that are not part of a cluster have been set to 0. **D)** The maximum contrast over the whole time course at 0 ms localized to the anterior part of the left putamen, and this peaked at 25 ms: The heat map is thresholded such that values that are not part of a cluster have been set to 0. **E)** and **F)** Time courses for the contrast between *Omission 0* and *Omission 15* with 95% confidence intervals across participants, extracted from the sources in **C** and **D. G)** and **H)** Time courses for *Omission 0, Omission 5* and *Omission 15*, extracted from the sources in **C** and **D**. Time courses in **E-H** are extracted from the source showing the maximum activation according to the automated anatomical labeling atlas (AAL) (Tzourio-Mazoyer et al., 2002).

In addition to the cerebellum and putamen, we also found differences in the thalamus, the inferior parietal cortex, SI and SII (Fig. 5), for the contrast between *Omission 0* and *Omission 15* in the beta band. In summary, we found that non-cortical structures, i.e. cerebellum, putamen and thalamus, followed the expected timing, i.e. the contrast peaking at ∼0 ms, whereas cortical structures such as SI and inferior parietal cortex peaked before the expected stimulus (∼-250 ms) and SII peaked after the expected stimulus (∼250 ms).

**Fig 5.**
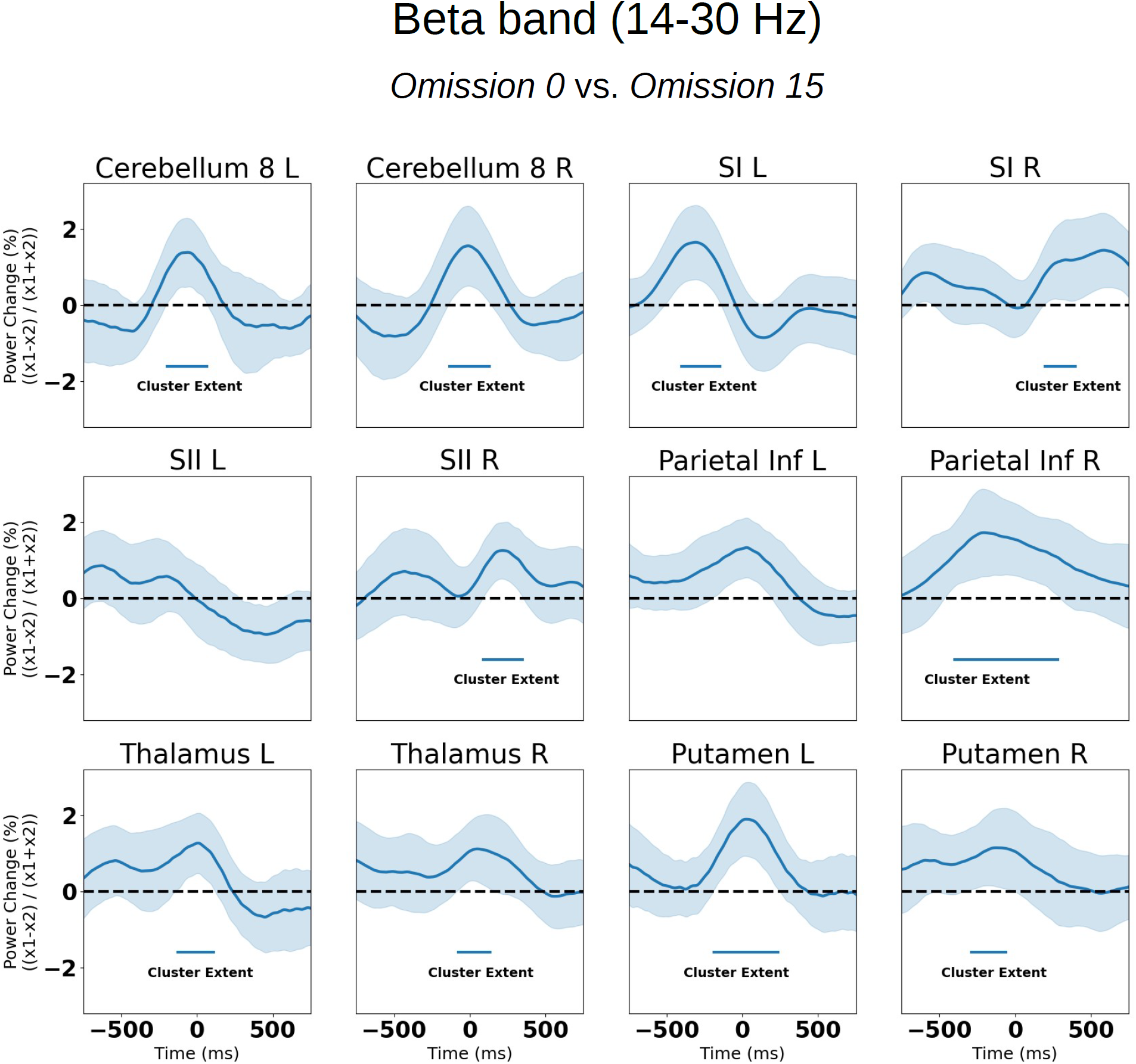
Omission contrast time courses for the areas of interest in the beta band: Time courses are extracted from the source showing the maximum contrast according to the automated anatomical labeling atlas (AAL) (Tzourio-Mazoyer et al., 2002) with the exception of SII, which is from the Harvard Oxford cortical atlas (Desikan et al., 2006). See Supplementary Fig. 1 for the individual time courses.

We then investigated the same areas for the contrast between *First Stimulation* and *Second Stimulation* for the theta band (Fig. 6). Here, we found differences in the very same areas. Except for the cerebellum, all differences were expressed around the peak of the left inferior parietal cortex (∼145 ms) (Fig. 3D&F&H). The cerebellar differences peaked around 285 ms for the left side and around 500 ms for the right side.

**Fig 6.**
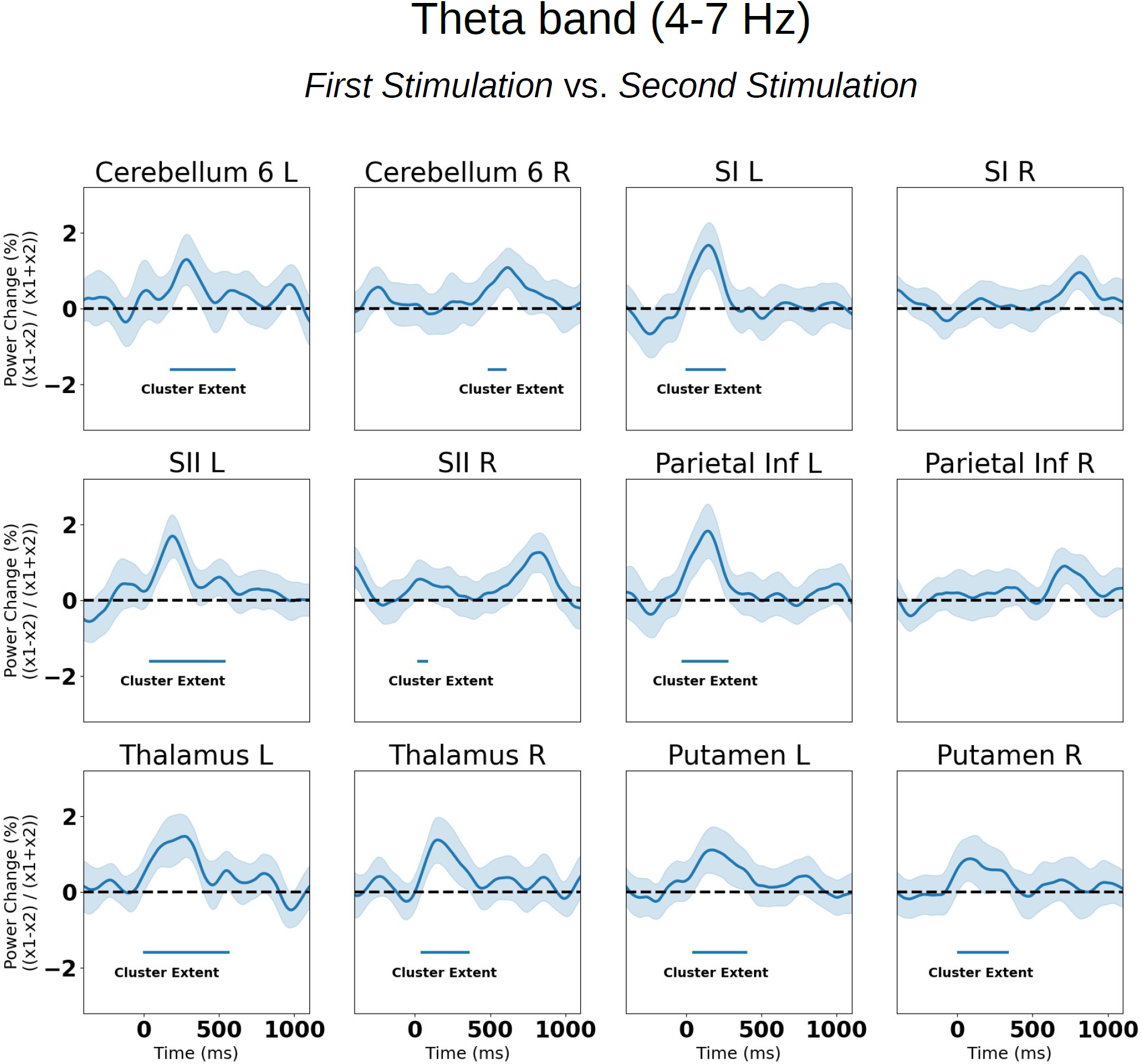
Stimulation contrast time courses for the areas of interest in the theta band: Time courses are extracted from the source showing the maximum contrast according to the automated anatomical labeling atlas (AAL) (Tzourio-Mazoyer et al., 2002) with the exception of SII, which is from the Harvard Oxford cortical atlas (Desikan et al., 2006). See Supplementary Fig. 2 for the individual time courses.

### 3.3 Control analyses

An alternative explanation for why we found differences in cerebellar activity when comparing omissions following regular and irregular trains of stimulation may be that the proposed cerebellar clock is based on local timelocking rather than global timelocking. By global timelocking, we mean that the timing of the expected, but omitted, stimulus is set to 8,982 ms (six times the ISI (1,497 ms)) after the first stimulation of the train (Fig. 1). By local timelocking, we mean that the timing of expected, but omitted, stimulus is set to the temporal distance between the fifth and sixth stimulation, whatever that may be in each particular stimulation train. Using local timelocking, we were also able to reject the null hypothesis (*p*_BIGGEST_CLUSTER_ = 0.0029), and we found similar activations of the cerebellum and the putamen (Supplementary Fig. 3). We also used mean timelocking, where we calculated mean distance between the last three stimulations (Fig. 1). Using mean timelocking, we found similar activations of the cerebellum and the putamen (Supplementary Fig. 4) (*p*_BIGGEST_CLUSTER_ = 0.0049). The way we timelocked the data in the main analysis (Fig. 4) thus does not change the results significantly. (For comparison, *p*_BIGGEST_CLUSTER_ = 0.0078, for the global timelocking reported in the main results).

It is also possible that the contributions of the gradient could not be filtered by the beamformer. We therefore re-ran the beamformer analysis on data where the signal space projections had been applied. This again resulted in very similar results (*Omission 0* vs. *Omission 15*; *p*_BIGGEST_CLUSTER_ = 0.0049; Supplementary Fig. 5).

## 4 Discussion

We found that our hypothesis, that cerebellum more strongly activates to omissions appearing in a regularly timed train than in an irregularly timed train, was corroborated by our beta band findings, showing that the contrast and the omission activity related to the regular train was at their strongest close to 0 ms (Fig. 4E & 4G).

### 4.1 A cerebellar clock (the beta band)

The differences in cerebellum are maximal around the expected time of stimulation, which indicates that cerebellar power reflects the strength of the temporal expectation. Cerebellar power thus reflects the prediction of an upcoming somatosensory stimulation, as beta band power is expected to increase when predicted information is expected to arrive (Engel and Fries, 2010) (Fig. 4E). The cerebellar findings here are also similar to findings from the auditory domain, where it has been shown that beta band power in the auditory cortex synchronizes with the rhythm of auditory stimuli, peaking at their occurrence and then decreasing again (Arnal and Giraud, 2012; Fujioka et al., 2012). Interestingly, the most irregular condition (*Omission 15*) showed a dip for oscillatory power around the time, the stimulation should have occurred, had the pattern been regular (Fig. 4G). In this condition, a stimulation is expected, but its exact timing is uncertain. Given that a function of prediction is to inhibit the processing of expected stimuli, we can interpret this as the cerebellum disinhibiting the processing of spatially expected, but temporally unexpected, stimuli.

Our results furthermore indicated that the putamen and thalamus are more strongly activated for the omission preceded by regular trains than omissions preceded by irregular trains. This fits well with the knowledge that the thalamus and basal ganglia are involved in timing as a recent meta-analysis of functional magnetic resonance imaging timing studies showed (Teghil et al., 2019). Of notice is that they follow the timing of the cerebellum, peaking around 0 ms, indicative of a network of cerebellum, putamen and thalamus (Bostan and Strick, 2018; Caligiore et al., 2016; Gibbon et al., 1997). For these, we also saw the dip in power for the most irregular condition (*Omission 15*) (Fig. 4H, Supplementary Fig. 1). Activations of putamen and thalamus found using MEG must be treated with caution due to their deep locations and the closed field of their neurons (Lorente de Nó, 1947). This results in a weak signal when measured outside the head that makes it challenging to say with confidence which deep source produced the MEG signal. However, theoretical work and simulation studies show that MEG should be sensitive to thalamus and (anterior) putamen (Attal et al., 2012; Attal and Schwartz, 2013). In terms of experimental evidence, it was recently shown that MEG can retrieve patterns of activation from the putamen and thalamus (Pizzo et al., 2019). This theoretical and experimental evidence together with the knowledge that the present paradigm should elicit activity in the basal ganglia and the thalamus combined with the high number of omission trials (*N*=150) mean that we believe that the present evidence is supportive, but not conclusive, of actual putamen and thalamus activations.

In contrast to the cerebellum, the SI and inferior parietal cortical contrasts peaked before (Fig. 5) the expected arrival of the stimulus and the SII contrast peaked after the expected arrival of the stimulus. The SI and inferior parietal cortex differences may indicate a dampening of cortical activity before the onset of an expected stimulus, compatible with the decrease in theta band activity for repeated stimulations (Figs. 3 & 6). For SI, this interpretation is supported by the findings of Shin et al. (2017) who find that failing to detect somatosensory stimuli correlates with an increase of beta activity before the stimulation and that attending away from a stimulus also increases beta power. In a similar manner, the function of the SI, and likewise that of inferior parietal cortex, beta activity may be to dampen the processing of an (expected) unsurprising stimulus. The SII activity may reflect an update of the current state of affairs, i.e. that the train of stimulation has been broken, and thus eliminating the prediction of a new stimulus. Similar results have been reported for omissions in the auditory domain, with beta band power increasing in the time interval after the expected, but omitted, stimulus in the auditory cortex (Fujioka et al., 2009).

In comparison to two recent studies (Andersen and Lundqvist, 2019; Naeije et al., 2018) we found weak evidence of an SII evoked response (Fig. 2). Both these studies used tactile stimulation by inflating a membrane fastened to the fingertips of participants and found an SII response around 140 ms time-locked to the expected, but omitted, stimulation. A possible explanation for not finding as strong evidence for such a response in the current study is that the instantaneousness of the electrical stimulation used in the current study relative to the longer extended touch of the membrane makes it harder for the brain to time-lock to the exact time point. Similarly, the older study of Tesche and Karhu (2000), using electrical stimulation of the median nerve, also only found weak evidence of an SII evoked response to the omissions. This would have the consequence that evoked analyses would be less sensitive to the SII response, when electrical stimulation is used as compared to mechanical stimulation.

To summarize, the current findings provide evidence that the cerebellum, together with the putamen and thalamus, track the timing of upcoming stimulation, as reflected by beta band power. This fits the interpretation of the beta band as predicting the *when* of upcoming stimulation (Arnal and Giraud, 2012), and supports the notion of the cerebellum as a clock that keeps track of upcoming stimulation. Cortical structures such as the SI, SII and the inferior parietal cortex are also timed relative to the upcoming stimulation. A potential explanation of the order of events is that SI and inferior parietal cortex increases in beta power indicate an internal attending away (Shin et al., 2017) from the expected, but omitted stimulus *before* it is expected to arrive. The cerebellar increase in power reflects the temporal tracking of the expected stimulus (Courchesne and Allen, 1997), with the SII activity indicating that the stimulus train has been broken, and that the current set of predictions is not valid anymore.

### 4.2 Encoding of expectations (theta band)

We furthermore found in the theta band that the cerebellum reacted more strongly to the first stimulation of a train compared to the subsequent stimulation around 280 ms after stimulation onset. This may reflect the unexpected nature of the first stimulation compared to the subsequent stimulation. This is similar to what has been found using electrical stimulation (Tesche and Karhu, 2000; see also: Naeije et al., 2018), but opposite to what has been found using tactile stimulation (Andersen and Lundqvist, 2019). Future studies will have to tell whether this difference is dependent on the type of stimulation. Similarly to the omission contrast, cortical areas, i.e. SI, SII, and inferior parietal cortex, and non-cortical areas, i.e., cerebellum, putamen and thalamus, were found to differ in activation between the first and the subsequent stimulation (Fig. 6). Except for the cerebellum, all contrasts peaked around 145 ms (Fig. 3F), whereas cerebellum peaked later, around 285 ms (Fig. 3E). All the cortical areas were expressed most strongly on the contralateral side of the stimulation as expected.. The later cerebellar activations are potentially an encoding of the new state of affairs (Engel and Fries, 2010), namely that a new stimulation sequence has begun. One might have expected the increase in cerebellar activity for the first stimulation relative to the second stimulation to be centred around 0 ms, if the cerebellum functions as a clock, tracking the stimuli. Leading up to a first stimulation, however, there is no prediction yet, as it has just been eliminated by the omission. It is possible that the cerebellar activation reflects an encoding of the new stimulation train, i.e. setting the clock, based on information from the earlier (145 ms) cortical and subcortical activations.

### 4.3 Cerebellum’s role in forming expectations

The current study reveals that cerebellum (along with its potential network members, thalamus and putamen) was the only area that peaked around 0 ms when comparing omissions (Fig. 5). We interpret this as the cerebellum clocking that stimuli arrive as expected, possibly in unison with the putamen and the thalamus (Bostan and Strick, 2018). However, a competing model (Merchant et al., 2013) has it that the core timing network is restricted to the basal ganglia and the thalamus, and that these core areas interact with specific structures depending on the context, e.g. for auditory timing, these areas would interact with the auditory cortex. In fact, such clocking activity has been found in the auditory cortex (Fujioka et al., 2012; Ruiz et al., 2017). According to this model, cerebellum is only involved in timing in contexts where motor output is required. Seemingly, this is at odds with the timing-related cerebellar activations also found by Fujioka et al. (2012) and Ruiz et al. (2017). However, this may be explained by the participants in these studies having to perform motor related tasks in response to stimuli. This would however not be able to explain that cerebellum is involved in timing in the current study, given that the task is purely somatosensory and no motor output is required. Our current results thus are compatible with the cerebellum being a core part of the timing network, as proposed by Bostan and Strick (2018). Lesion studies also indicate this (Ivry, 1993; Ivry and Keele, 1989) with cerebellar patients showing deficiencies in both perception and production in timing tasks. At a more general level, the present findings may be encompassed by the theory of the cerebellum as having a predictive and a preparatory function (Courchesne and Allen, 1997), setting the internal conditions for the mental or motor output that a given context may require. Part of this predictive activity would then be clocking when stimuli should arrive. Our results suggest that this clocking activity is expressed in the beta band.

We also found cerebellar activity in the theta band when comparing the first stimulation to the subsequent stimulation. In contrast to the beta band activity, the theta cerebellar activity was unique in peaking later (∼285 ms) than the first stimulation related activations (∼145 ms) (Fig. 6). We interpret this as the cerebellum updating its clock for subsequent stimulation. Of note is also that, in contrast to the omission-related beta band activity, the putamen and the thalamus peak out of sync with the cerebellum. Our findings are similar to recent findings (Dave et al., 2020), where transcranial magnetic stimulation was used to knock out cerebellar theta and cerebellar beta respectively. Cerebellar theta was found to be related to encoding episodic memories and cerebellar beta was found to be related to semantic predictions related to those memories.

The present findings in the theta and beta band ranges fit well with the fact that the granular layer of the cerebellum supports low-frequency oscillations (D’Angelo et al., 2009). However, given that the only output of the cerebellar cortex comes from the Purkinje cells in the deep cerebellar nuclei, one might expect to find high-frequency oscillations in the 160-200 Hz range, as have been found in local field potential recordings in mice and humans (Cheron et al., 2008; Dalal et al., 2013). Oscillations this fast may make possible the temporal precision on the order of tens of milliseconds found here, and may thus play an important role as well. Reports of high-frequency cerebellar oscillations using non-invasive recordings are scarce however (but see: Todd et al., 2018), likely due to the very low signal-to-noise ratio for high frequencies but on-scalp MEG (Boto et al., 2019; Pfeiffer et al., 2020) may be sensitive to these high-frequency oscillations.

Furthermore, we find intriguing evidence of putamen and thalamus activation, tracking the timing of somatosensory stimuli as well. This fits well with knowledge from other domains, such as functional magnetic resonance imaging. However, given the low sensitivity of magnetoencephalography to the putamen and the thalamus, further research is warranted.

In conclusion, we find that the cerebellum functions as a clock that tracks rhythmic stimuli. We interpret the cerebellum’s theta oscillations as establishing the proposed cerebellar clock, while the cerebellum’s beta activity reflects comparison of incoming stimuli with the clock’s prediction.

## 5 Acknowledgements

We thank Daniel Lundqvist for providing the lab equipment at the National Facility for Magnetoencephalography (NatMEG) at Karolinska Institutet for the very first pilots. We thank Johannes Singer, who was invaluable in designing and analysing pilot data, leading to the final design. Furthermore, we thank Sigbjørn Hokland and Marie-Louise Holm Møller for their help with data collection. Lau Møller Andersen was funded by the Carlsberg Foundation (CF18-0843) and by the Aarhus University Research Foundation (AUFF-E-2019-9-20). Sarang Dalal was funded by an European Research Council Starting Grant (640448).

