## Supplementary Figures 1-5 for "The cerebellar clock: predicting and timing somatosensory touch"

### Beta band (14-30 Hz)

*Omission 0* vs. *Omission 15*

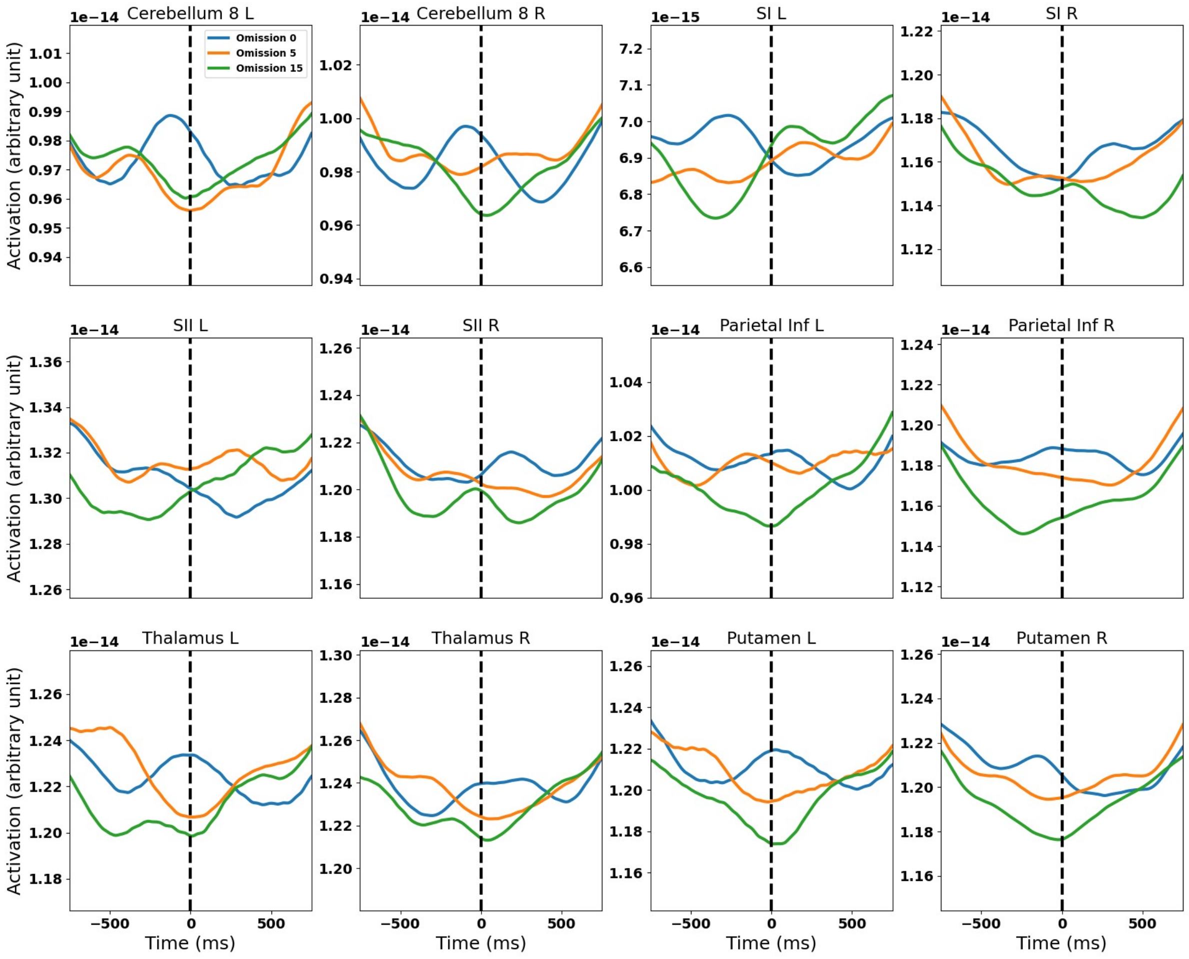


**Supplementary Fig. 1: Omission time courses for the areas of interest in the beta band:** Time courses are extracted from the source showing the maximum contrast according to the automated anatomical labeling atlas (AAL) (Tzourio-Mazoyer et al., 2002) with the exception of SII, which is from the Harvard Oxford cortical atlas (Desikan et al., 2006). See Fig. 5 for the contrast between *Omission 0* and *Omission 15*.

### Theta band (4-7 Hz)

*First Stimulation* vs. *Second Stimulation*

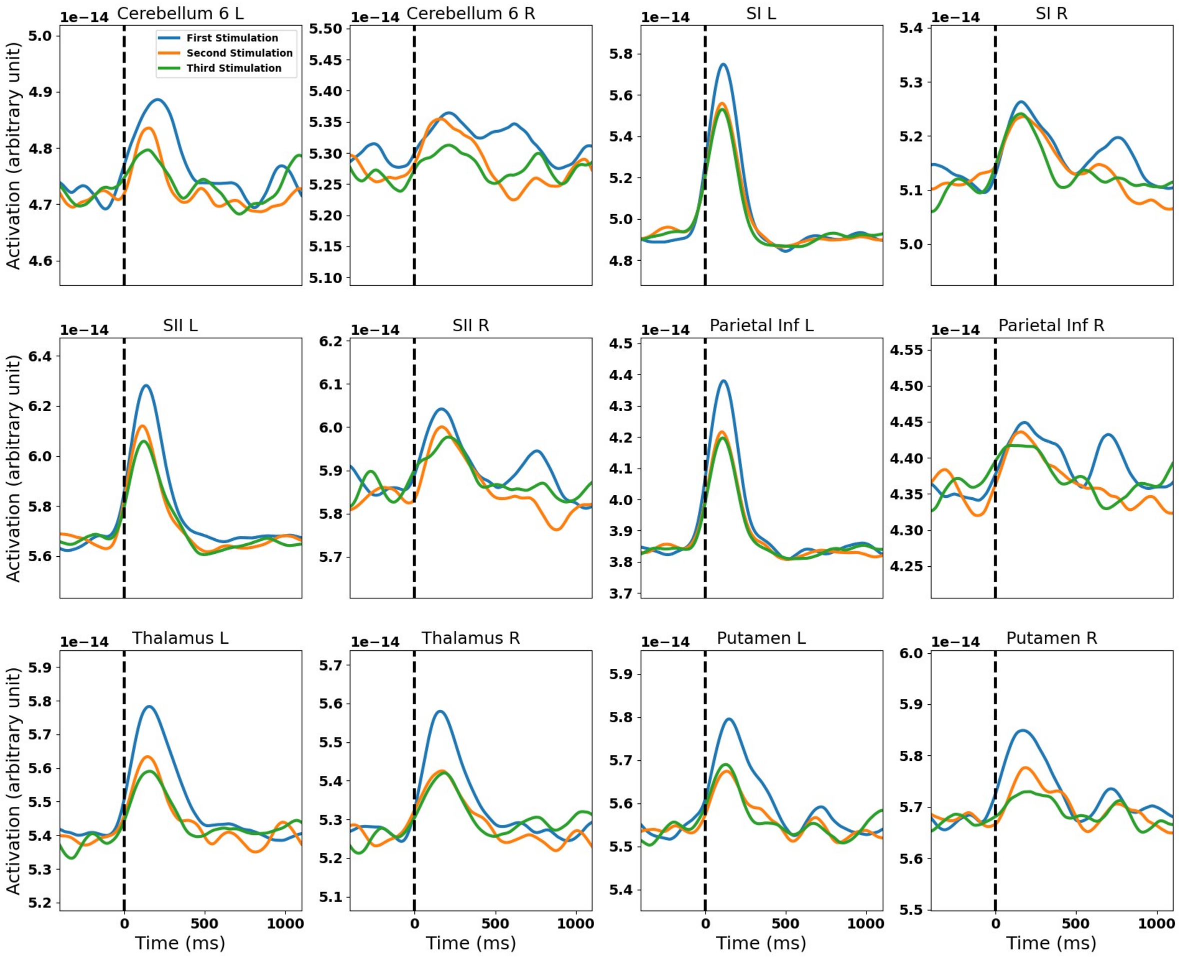


**Supplementary Fig. 2: Stimulation time courses for the areas of interest in the theta band:** Time courses are extracted from the source showing the maximum contrast according to the automated anatomical labeling atlas (AAL) (Tzourio-Mazoyer et al., 2002) with the exception of SII, which is from the Harvard Oxford cortical atlas (Desikan et al., 2006). See Fig. 6 for the contrast between *First Stimulation* and *Repeated Stimulation (2)*.

### Beta band (14-30 Hz)

#### *Omission 0* vs. *Omission 15*

#### *Local timelocking*

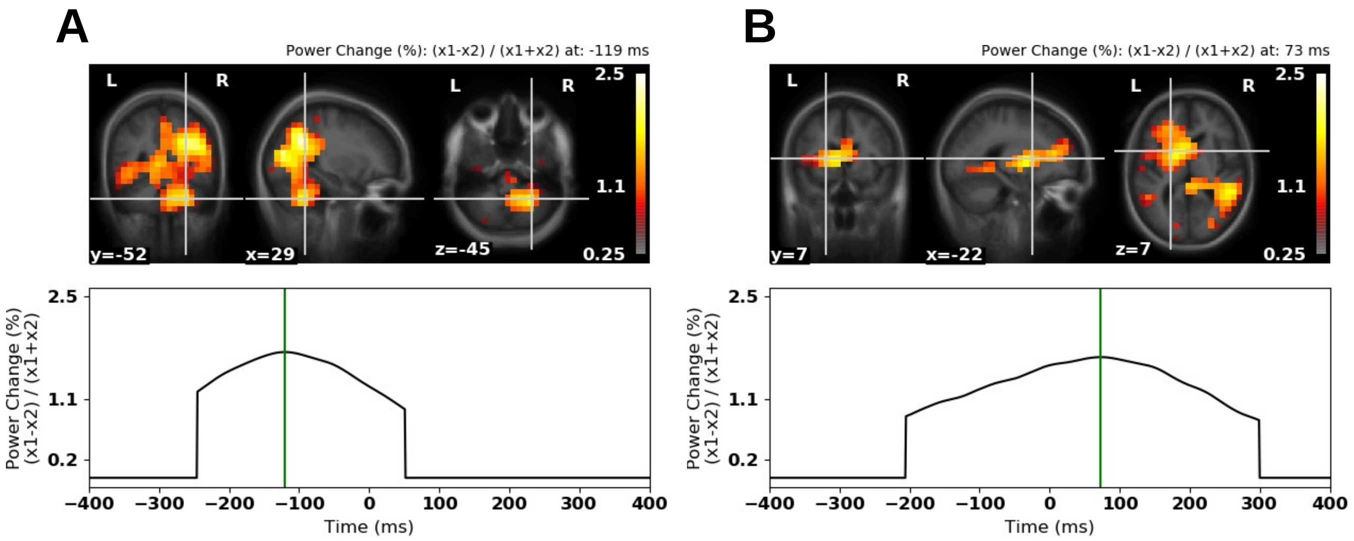


**Supplementary Fig. 3: Differences between *Omission 0* and *Omission 15* in the beta band (14-30 Hz) (-400 ms to 400 ms) using local timelocking A)** Contrast localized to the right cerebellum at -119 ms: Both the heat map and the time course below are thresholded such that values that are not part of a cluster have been set to 0. **B)** Contrast localized to the left putamen at 73 ms: Both the heat map and the time course below are thresholded such that values that are not part of a cluster have been set to 0. The contrast value expresses the difference in power as a percentage of total power in the two conditions. For both positions, the source with the maximum value in the original analysis was chosen.

### Beta band (14-30 Hz)

*Omission 0* vs. *Omission 15*

*Mean timelocking*

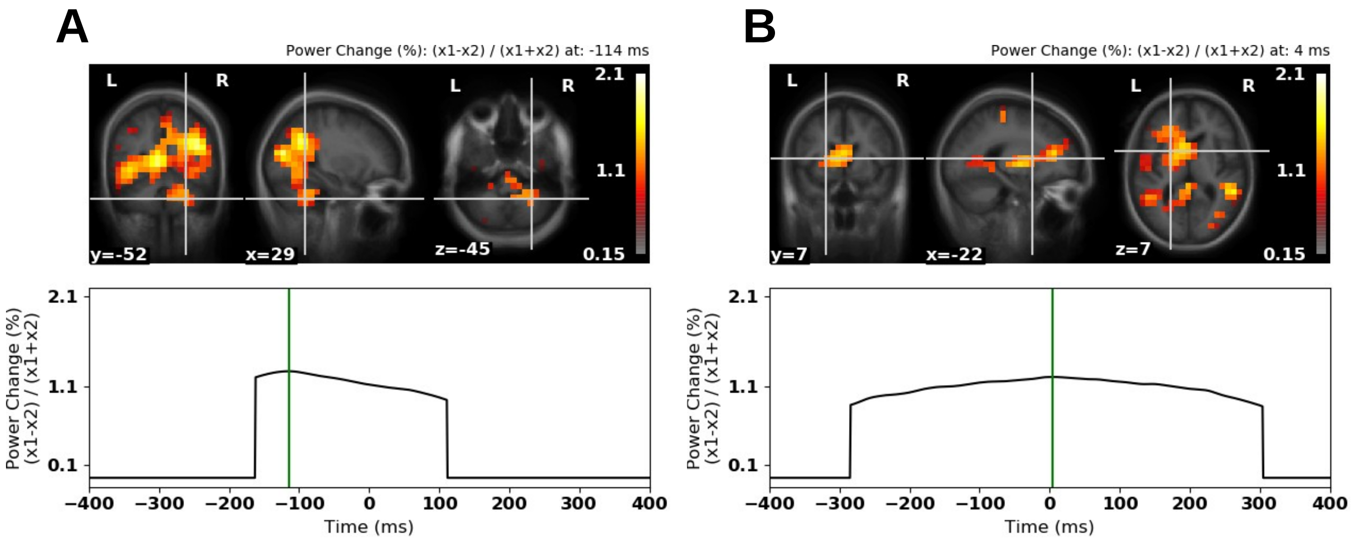


**Supplementary Fig. 4: Differences between *Omission 0* and *Omission 15* in the beta band (14-30 Hz) (-400 ms to 400 ms) using mean timelocking A)** Contrast localized to the right cerebellum at -114 ms: Both the heat map and the time course below are thresholded such that values that are not part of a cluster have been set to 0. **B)** Contrast localized to the left putamen at 4 ms: Both the heat map and the time course below are thresholded such that values that are not part of a cluster have been set to 0. The contrast value expresses the difference in power as a percentage of total power in the two conditions. For both positions, the source with the maximum value in the original analysis was chosen.

### Beta band (14-30 Hz)

*Omission 0* vs. *Omission 15*

*Projections applied*

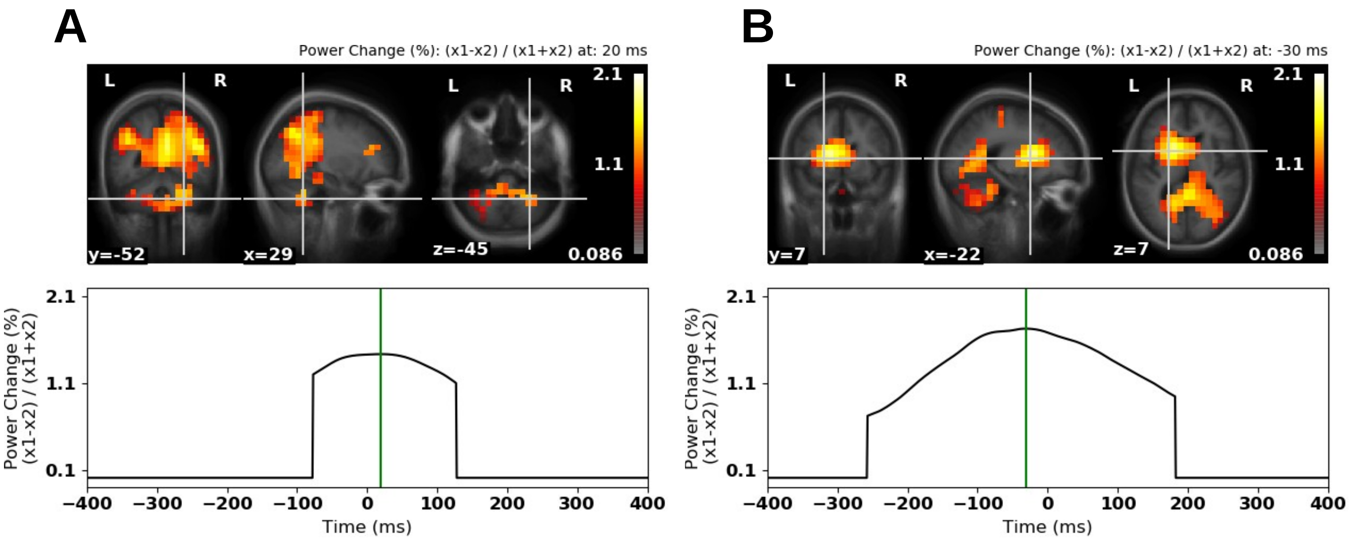


**Supplementary Fig. 5: Differences between *Omission 0* and *Omission 15* in the beta band (14-30 Hz) (-400 ms to 400 ms) with signal space projections applied A)** Contrast localized to the right cerebellum at 20 ms: Both the heat map and the time course below are thresholded such that values that are not part of a cluster have been set to 0. **B)** Contrast localized to the left putamen at -30 ms: Both the heat map and the time course below are thresholded such that values that are not part of a cluster have been set to 0. The contrast value expresses the difference in power as a percentage of total power in the two conditions. For both positions, the source with the maximum value in the original analysis was chosen.
