## Supplementary Tables 1-2 for "The cerebellar clock: predicting and timing somatosensory touch"

| Region | Begin time | End time |
|---|---|---|
| Insula_L | -89 ms | 48 ms |
| Cingulum_Mid_L | 203 ms | 400 ms |
| Cingulum_Mid_R | 214 ms | 400 ms |
| Cingulum_Post_L | -46 ms | 54 ms |
| Cingulum_Post_R | -146 ms | 135 ms |
| Lingual_L | -91 ms | 43 ms |
| Parietal_Sup_R | -28 ms | 57 ms |
| Parietal_Inf_R | -19 ms | 278 ms |
| Supramarginal_R | 195 ms | 229 ms |
| Angular_R | -395 ms | 199 ms |
| Precuneus_R | -78 ms | 85 ms |
| Paracentral_Lobule_L | -34 ms | 354 ms |
| Paracentral_Lobule_R | -28 ms | 400 ms |
| Caudate_L | -123 ms | 235 ms |
| Putamen_L | -160 ms | 159 ms |
| Pallidum_L | -127 ms | 173 ms |
| Temporal_Inf_L | -228 ms | 48 ms |
| Cerebelum_8_R | -59 ms | 11 ms |
| Cerebelum_9_R | -60 ms | 141 ms |

**Table 1: Regions part of a cluster for Omission 0 vs Omission 15 using the automated anatomical labeling atlas (Tzourio-Mazoyer et al., 2002):** The time ranges include only time points where at least 50 % of the voxels in the given region are part of a cluster. It is extremely important to note that the time ranges are included only as a *descriptive statistic* – the precise range cannot be *inferred* from a cluster permutation test (Sassenhagen and Draschkow, 2019).

| Region | Begin time | End time |
|---|---|---|
| Precentral_L | 47 ms | 257 ms |
| Frontal_Sup_Orb_L | 24 ms | 186 ms |
| Frontal_Sup_Orb_R | 271 ms | 354 ms |
| Frontal_Mid_L | 123 ms | 246 ms |
| Frontal_Mid_Orb_L | 21 ms | 166 ms |
| Frontal_Inf_Oper_l | 75 ms | 302 ms |
| Frontal_Inf_Oper_R | 333 ms | 361 ms |
| Frontal_Inf_Tri_L | 47 ms | 287 ms |
| Frontal_Inf_Orb_L | 66 ms | 212 ms |
| Frontal_Inf_Orb_R | 275 ms | 322 ms |
| Rolandic_Oper_L | 90 ms | 322 ms |
| Rolandic_Oper_R | 69 ms | 194 ms |
| Frontal_Med_Orb_L | 139 ms | 194 ms |
| Frontal_Med_Orb_R | 263 ms | 391 ms |
| Insula_L | 183 ms | 278 ms |
| Cingulum_Ant_R | 341 ms | 380 ms |
| Cingulum_Mid_L | 437 ms | 465 ms |
| Cingulum_Post_L | 33 ms | 163 ms |
| Cingulum_Post_R | 309 ms | 495 ms |
| ParaHippocampal_L | 271 ms | 292 ms |
| Calcarine_R | 477 ms | 600 ms |
| Cuneus_L | 86 ms | 100 ms |

| | | |
|---|---|---|
| Lingual_R | 492 ms | 600 ms |
| Occipital_Mid_R | 515 ms | 600 ms |
| Occipital_Inf_R | 316 ms | 355 ms |
| Fusiform_L | 190 ms | 327 ms |
| Fusiform_R | 586 ms | 600 ms |
| Postcentral_L | 71 ms | 266 ms |
| Parietal_Sup_L | -35 ms | 350 ms |
| Parietal_Inf_L | -20 ms | 283 ms |
| Supramarginal_L | 56 ms | 315 ms |
| Angular_L | 158 ms | 215 ms |
| Caudate_L | 139 ms | 343 ms |
| Putamen_L | 191 ms | 307 ms |
| Putamen_R | 310 ms | 326 ms |
| Pallidum_L | 235 ms | 325 ms |
| Thalamus_L | 54 ms | 373 ms |
| Thalamus_R | 69 ms | 346 ms |
| Heschl_L | 105 ms | 298 ms |
| Temporal_Sup_L | 70 ms | 298 ms |
| Temporal_Pole_Sup_L | 178 ms | 193 ms |
| Temporal_Mid_L | 102 ms | 297 ms |
| Temporal_Inf_L | 228 ms | 316 ms |
| Cerebelum_4_5_L | 269 ms | 273 ms |
| Cerebelum_4_5_R | 534 ms | 600 ms |
| Cerebelum_6_L | 206 ms | 291 ms |
| Cerebelum_6_R | 539 ms | 600 ms |
| Cerebelum_8_L | 77 ms | 99 ms |
| Cerebelum_8_R | 550 ms | 600 ms |
| Cerebelum_9_L | 84 ms | 600 ms |
| Cerebelum_9_R | 492 ms | 596 ms |
| Vermis_4_5 | 473 ms | 508 ms |
| Vermis_9 | 554 ms | 600 ms |
| Vermis_10 | 495 ms | 600 ms |

**Table 2: Regions part of a cluster for First Stimulation vs Repeated Stimulation (2) using the automated anatomical labeling atlas (Tzourio-Mazoyer et al., 2002)::** The time ranges include only time points where at least 50 % of the voxels in the given region are part of a cluster. It is extremely important to note that the time ranges are included only as a *descriptive statistic* – the precise range cannot be *inferred* from a cluster permutation test (Sassenhagen and Draschkow, 2019).
